# Modelling CAR T-cell Therapy with Patient Preconditioning

**DOI:** 10.1101/2020.06.20.162925

**Authors:** Katherine Owens, Ivana Bozic

## Abstract

The Federal Drug Administration (FDA) approved the first Chimeric Antigen Receptor T-cell (CAR T-cell) therapies for the treatment of several blood cancers in 2017, and efforts are underway to broaden CAR T technology to address other cancer types. Standard treatment protocols incorporate a preconditioning regimen of lymphodepleting chemotherapy prior to CAR T-cell infusion. However, the connection between preconditioning regimens and patient outcomes is still not fully understood. Optimizing patient preconditioning plans and reducing the CAR T-cell dose necessary for achieving remission could make therapy safer. In this paper, we test treatment regimens consisting of sequential administration of chemotherapy and CAR T-cell therapy on a system of differential equations that models the tumor-immune interaction. We use numerical simulations of treatment plans from within the scope of current medical practice to assess the effect of preconditioning plans on the success of CAR T-cell therapy. Model results affirm clinical observations that preconditioning can be crucial for some patients, not just to reduce side effects, but to even achieve remission at all. We demonstrate that preconditioning plans using the same CAR T-cell dose and the same total concentration of chemotherapy can lead to different patient outcomes due to different delivery schedules. Results from sensitivity analysis of the model parameters suggest that making small improvements in the effectiveness of CAR T-cells in attacking cancer cells, rather than targeting the recruitment and longevity of CAR T-cells, will significantly reduce the minimum dose required for successful treatment. Our modeling framework represents a starting point for evaluating the efficacy of patient preconditioning in the context of CAR T-cell therapy.

## 1 Introduction

### 1.1 CAR T-cell therapy

Cancer treatment plans often draw upon multiple treatment modalities, including surgery, radiation, chemotherapy, and immunotherapy, in order to combat disease (Miller et al., 2019; Mokhtari et al., 2017; Palmer and Sorger, 2017; Khalil et al., 2016; Yamamoto et al., 2016). Immunotherapies aim to enhance the body’s natural defense mechanisms against tumor cells. Immunotherapy approaches include injections of molecules that promote immune activity like interleukins, antibodies, checkpoint inhibitors, and vaccines, as well as adoptive cellular therapies (ACT) in which T-cells are isolated from the patient, expanded ex vivo, and then reintroduced into the patient (Kruger et al., 2019).

Advances in gene-editing during the 1990s enabled a new type of ACT in which a patient’s T-cells are genetically engineered to recognize markers displayed on the surface of tumor cells (Almåsbak et al., 2016). These genetically engineered T-cells, called CAR T-cells, were first reported to effectively eradicate lymphoma cells in a murine model in 2010 (Kochenderfer et al., 2010). Following this breakthrough, CAR T-cell therapy showed dramatic success against lymphomas and leukemias in a series of clinical trials with complete remission rates in the range of 65-90% (Almåsbak et al., 2016). In late 2017 the FDA approved two CAR T-cell therapies for treatment of relapsed and refractory Acute Lymphoblastic Leukemia and for diffuse large B-cell lymphoma (June and Sadelain, 2018). Continued advances in T-cell engineering, genetic editing, selection of targets, and cell manufacturing have the potential to broaden applications of CAR T-cell therapy to treat diseases beyond leukemia and lymphoma (Almåsbak et al., 2016; Brown and Mackall, 2019; Pettitt et al., 2018).

Although CAR T-cell therapies show great promise in the fight against cancer, there are multiple challenges hindering their universal adoption, including potentially fatal inflammatory side effects that occur in up to 10% of patients who have positive responses to the infusion (Pettitt et al., 2018). The toxic side effects of CAR T-cell therapy include neurologic effects, B-cell aplasia, and Cytokine Release Syndrome (CRS), ranging in severity from mild flu-like symptoms to death. Studies associate higher levels of tumor burden at the time of treatment with more serious side effects (Brown and Mackall, 2019).

Under current medical practice, care providers administer a lymphodepleting round of chemotherapy prior to CAR T-cell injections in order to increase efficacy and reduce side effects of treatment (Mahadeo et al., 2019). It has been hypothesized that by reducing tumor burden and the number of normal immune cells, chemotherapy allows CAR T-cells to proliferate and overcome the cancer cells more easily. Though the use of such preconditioning regimen is common, the optimal combination of chemotherapy and CAR T-cells has not yet been determined. A mathematical model illustrating how these two forms of treatment interact for any given patient could help address this question.

### 1.2 Mathematical modeling of cancer treatment

Systems of ordinary differential equations (ODEs) have been widely used to model tumor-immune interactions on the cell population level (Borges et al., 2014; de Pillis et al., 2005, 2006; Kirschner and Panetta, 1998; Kuznetsov et al., 1994; Nanda et al., 2013; Pinho et al., 2002; Ribba et al., 2012; Rösch et al., 2016; Usher, 1994). Many of these models also incorporate treatment via chemotherapy, immunotherapy or both. Several thorough reviews of non-spatial mathematical models of interactions between the immune system, tumor cells, and cancer therapies exist in the literature (Eftimie et al., 2011; Talkington et al., 2018).

Methods used to mathematically model cancer and the immune response range in complexity from using two differential equations to describe predator-prey type dynamics between immune and tumor cells (Usher, 1994; Kirschner and Panetta, 1998; Kuznetsov et al., 1994; Sahoo et al., 2020; Talkington et al., 2018), up through detailed mechanistic models that explicitly represent multiple components of the immune response (Hardiansyah and Ng, 2019; Harris et al., 2018; Ribba et al., 2012). Cancer therapy has been incorporated through simple time-independent dosing (Usher, 1994; Kirschner and Panetta, 1998; Pinho et al., 2002), as well as through more realistic time-dependent dose schedules (Pinho et al., 2002; de Pillis et al., 2006). Here we aim to formulate a goldilocks model, incorporating just enough complexity to achieve expected biological outcomes and enable testing a range of medically relevant treatment plans (Brady and Enderling, 2019).

Several computational models have recently been developed to investigate aspects of CAR T-cell therapy including cytokine release syndrome toxicity management (Hopkins et al., 2018; Stein et al., 2017, 2019), mechanisms of CAR T-cell activation (Harris et al., 2018; Rohrs et al., 2019), and how factors including CAR T-cell dose, donor-dependent T-cell differences, cancer cell proliferation, and target antigen expression contribute to the overall effectiveness of CAR T-cell therapy (Hardiansyah and Ng, 2019; Kimmel et al., 2019; Rodrigues et al., 2019; Sahoo et al., 2020). Here we formulate a model that allows us to investigate how the interplay between specific preconditioning plans and CAR T-cell dosage affects patient outcomes.

The remainder of the paper follows this general outline: In Section 2, we first formulate and analyze the model for tumor-immune system interaction in the absence of treatment. Then, we detail how sequential combination of chemotherapy and CAR T-cell therapy is incorporated into the model. In Section 3 we evaluate numerical simulations of treatment plans that adhere to the FDA package inserts for the two approved CAR T-cell therapies (Kite, 2017; Novartis, 2017). As discussed in Section 4, these simulations show that appropriate chemotherapeutic preconditioning can reduce the CAR T-cell dosage necessary for a patient to achieve remission. Our results suggest that small changes in the delivery schedule of chemotherapy and CAR T-cell therapy can have a significant effect on the outcome of treatment.

## 2 Model

### 2.1 Model without treatment

Here we formulate a model of the tumor-immune interaction by updating Kuznetsov’s influential 1994 model (Kuznetsov et al., 1994). Our goal is to incorporate experimentally validated interaction terms while maintaining the simplicity of low-dimensional models of tumor-immune interaction. In order to make useful suggestions about potential treatment plans, we require our model to meet reasonable biological assumptions. With appropriate parameter values, we expect to observe

- Uncontrolled growth of tumor cells in the absence of immune response,
- Immune cells reduce the number of tumor cells,
- Both immune cells and tumor cells decrease under chemotherapy.

Additionally, we want the model to reflect that additional immune cells are activated and recruited by interactions between cancer cells and immune cells. The activation of tumor-killing cytotoxic T-cells is dependent on interactions between surface molecules on the T-cell and a tumor cell. Once activated, a T-cell undergoes clonal expansion, increasing the number of cells specific for the target antigen that can then travel throughout the body in search of antigen-positive tumor cells (Harlin et al., 2009; Smith-Garvin et al., 2009). Additional T-cells may also be activated by fragments of tumor cells that have been lysed by other cells (Huang et al., 1994).

Finally, we want our model to reflect that immune cells can become inactivated through repeated interaction with tumor cells. Upon repeated exposure to tumor cells some T-cells experience exhaustion, an altered functional state in which they cannot effectively attack foreign cells (Thommen and Schumacher, 2018).

We started from Kuznetsov’s model, which includes only one group of activated immune cells or “effector” cells rather than multiple subpopulations of immune cells (Kuznetsov et al., 1994). Kuznetsov’s model is

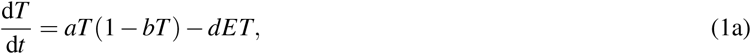

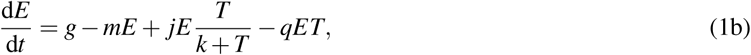

where *T* (*t*) is the population of tumor cells at time *t* and *E*(*t*) is the population of effector cells at time *t*. For our model, we replace the second term in Eq. (1a) and the third term in Eq. (1b) with experimentally-validated, ratio-dependent interaction terms which de Pillis et al. (2005) demonstrated achieve a better fit to experimental data from cytotoxicity assays compared to simpler, classical choices such as mass action or Michaelis-Menten (which are used in the Kusnetsov model). We chose to adopt their novel interaction terms into the Kuznetsov model, rather than working with de Pillis et al.’s 5-equation model system, in order to focus on the minimal set of equations that would produce desired biological behaviors.

In sum, our model for tumor-immune interation in the absence of treatment is

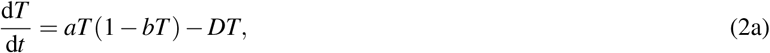

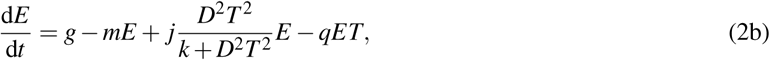

where *D* describes the tumor-cell lysis rate (rate at which tumor cells are killed by effector cells)

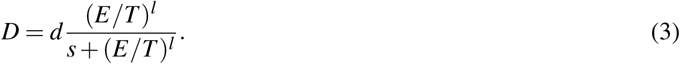

In the absence of effector cells, tumor cells grow logistically towards a carrying capacity of *b*^−1^, and in the absence of tumor cells, effector cells are produced at a fixed rate, *g*, and decay at a rate proportional to the population, *mE*. Effector cells kill tumor cells according to the ratio-dependent tumor cell death term, *DT*. Effector cells are recruited at a rate dependent on tumor cell lysis, *jED*^2^*T* ^2^*/*(*k* + *D*^2^*T* ^2^). The inactivation of effector cells upon exposure to tumor cells is incorporated through a mass-action term, *qET*, capturing the phenomenon of immune-cell exhaustion (Thommen and Schumacher, 2018). Model parameters are defined in Table 1, and for details of parameter selection, please refer to Appendix B.

**Table 1:** Parameter values from ranges for large B-cell lymphoma reported by Rösch et al. (2016) were selected for Patient 1. Parameter values from ranges for B cell Chronic Lymphocytic Leukemia reported by Nanda et al. (2013) were selected for Patient 2. The Patient 3 and 4 parameter values come from two metastatic melanoma patients, reported by de Pillis et al. (2005).

| Parameter | Description | Patient 1 | Patient 2 | Patient 3 | Patient 4 | Range |
| --- | --- | --- | --- | --- | --- | --- |
| a | tumor growth rate ($\text{day}^{-1}$) | $2.55\text{e}-1$ | $1.76\text{e}-1$ | $5.14\text{e}-1$ | $5.14\text{e}-1$ | $[4.5\text{e}-1, 5.5\text{e}-1]$ |
| b | 1/b is the carrying capacity (cells) | $1\text{e}-13$ | $1\text{e}-13$ | $1.02\text{e}-9$ | $1.02\text{e}-9$ | $[1\text{e}-13, 1\text{e}-8]$ |
| d | saturation level of fractional tumor kill by effector cells ($\text{day}^{-1}$) | 2.55 | 3.4 | 5.8 | 4.23 | $[1.5, 6]$ |
| g | base recruitment rate of effector cells (cells/day) | $1.4\text{e}3$ | $4.7\text{e}4$ | $1.3\text{e}4$ | $1.3\text{e}4$ | $[1\text{e}3, 5\text{e}4]$ |
| j | max recruitment rate of effector cells by tumor lysis ($\text{day}^{-1}$) | $1.1\text{e}-1$ | $4.46\text{e}-3$ | $3.75\text{e}-2$ | $3.75\text{e}-2$ | $[1\text{e}-3, 1\text{e}-1]$ |
| k | steepness of effector cell recruitment curve ($\text{cells}^2\text{day}^{-2}$) | $2.019\text{e}5$ | $7.0\text{e}7$ | $2.0\text{e}7$ | $2.0\text{e}7$ | $[1\text{e}7, 6\text{e}7]$ |
| l | exponent of fractional tumor cell kill by effector cells (unit-less) | 1.395 | 2 | 1.36 | 1.43 | $[1, 2.5]$ |
| m | death rate of effector cells ($\text{day}^{-1}$) | $7\text{e}-3$ | $3.4\text{e}-2$ | $2\text{e}-2$ | $2\text{e}-2$ | $[1\text{e}-2, 10]$ |
| q | inactivation of effector cells by tumor ($\text{cells}^{-1}\text{day}^{-1}$) | $3.42\text{e}-10$ | $1.43\text{e}-8$ | $3.42\text{e}-10$ | $3.42\text{e}-10$ | $[1\text{e}-10, 1\text{e}-6]$ |
| s | steepness of fractional tumor cell kill by effector cells (unit-less) | $3.05\text{e}-1$ | $3.05\text{e}-1$ | $2.5\text{e}-1$ | $3.6\text{e}-1$ | $[1\text{e}-2, 5\text{e}-1]$ |
| $K_T$ | fractional tumor cell kill by chemotherapy ($\text{day}^{-1}$) | | | | | $9.00\text{e}-1$ |
| $K_E$ | fractional healthy cell kill by chemotherapy ($\text{day}^{-1}$) | | | | | $6.00\text{e}-1$ |
| $\gamma$ | chemotherapy decay rate ($\text{day}^{-1}$) | | | | | $9.00\text{e}-1$ |

We show analytically that for a wide range of biologically relevant parameters, the system has two stable equilibria. A zero-tumor (or tumor-free) equilibrium occurs at *T* = 0 and *E* = *g/m*, and a high-tumor equilibrium occurs when the tumor cell population is near the carrying capacity, *b*^−1^, and there is a low number of effector cells. Nullclines and equilibria are plotted in Fig. 1a for the non-dimensionalized system, in which case the tumor-free equilibrium is at the point (*x* = 0, *y* = 1*/m*) and the high-tumor equilibrium is at the point (*x* ≈ 1, *y* ≈ 1*/*(*q* + *m*)). Trajectories that approach the zero-tumor equilibrium are considered to have a “healthy” outcome, whereas trajectories that approach the high-tumor equilibrium are considered to have an “unhealthy” outcome.

**Figure 1:**
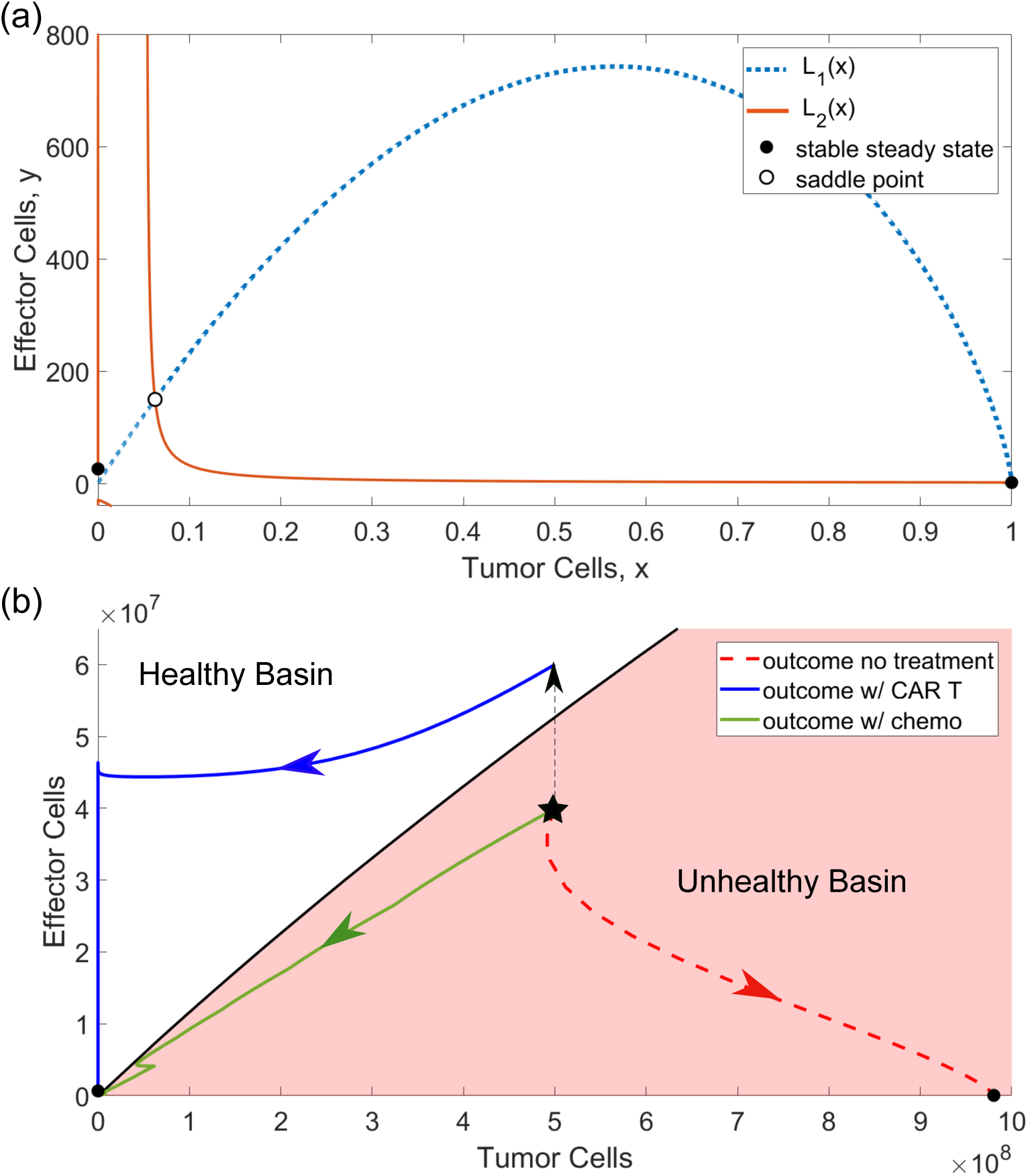
(a) Nullclines of the non-dimensionalized system without treatment. The two curves, *L*_1_(*x*) and *L*_2_(*x*), whose intersections determine the non-zero-tumor equilibrium are plotted for parameters from column 3 of Table 1. Circles mark the stable high-tumor steady state for *x* just less than 1 and the stable tumor-free steady state at the intersection between *L*_2_(*x*) and *x* = 0. There is also a saddle point where *L*_1_(*x*) and *L*_2_(*x*) intersect near a vertical asymptote of *L*_2_(*x*), indicated by a crossed circle. (b) Using parameters from column 4 of Table 1, the phase plane is partitioned into the basin of attraction for the high-tumor equilibrium (shaded) and the basin for the zero tumor equilibrium (white), with stable fixed points marked by circles. The goal of treatment is to shift a patient’s condition from the unhealthy basin of attraction to the healthy basin of attraction. Observe that without treatment, a trajectory starting at the starred initial condition approaches the high-tumor equilibrium over time. CAR T cell injections shift the patient into the healthy basin by increasing the number of effector cells, after which the trajectory approaches the tumor-free equilibrium. Alternatively, with a chemotherapy regimen, the patient is shifted into the healthy basin by decreasing the number of tumor cells. The resulting trajectory crosses the boundary between the basins at 5.×35 10^5^ tumor cells and 3. .× 98 10^4^ effector cells before approaching the tumor-free equilibrium

We also determined numerically that, as indicated in Fig. 1a, there is a saddle point for a relatively small number of tumor and effector cells. The stable manifold of this saddle point forms the separatrix between the basins of attraction for the zero-tumor and high-tumor equilibria (Fig. 1b). Note that in Fig. 1b we are only plotting the positive quadrant because it is an invariant set for this system. Thus, if we start with a non-negative initial condition, the model will never predict a negative population of cells, a desired property for biological interpretability. For the details of model analysis, please refer to Appendix A.

### 2.2 Model with treatment

The goal of treatment is to shift an initial patient condition from within the basin of attraction of the high-tumor equilibrium (unhealthy patient outcome) to the basin of attraction for the zero-tumor equilibrium (healthy patient outcome) through an appropriate intervention term. We consider intervention via chemotherapy and CAR T-cell injections. As illustrated in Fig. 1b, CAR T-cell injections shift the initial condition into the healthy basin by increasing the number of effector cells while chemotherapy shifts a trajectory into the healthy basin by decreasing the number of tumor cells.

Let *C*(*t*) be the concentration of a chemotherapy drug at time *t*. Following the method of de Pillis et al. (2006), we introduce a governing equation for the drug concentration

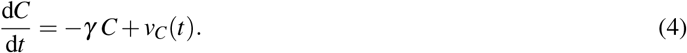

Here *γ* is the decay rate of chemotherapy and *v*_*C*_(*t*) is a time-dependent forcing term encoding the dose strength, *S* (*µ*M/day). The dosing schedule is

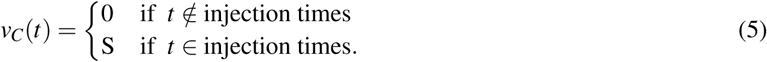

On the inset axis of Fig. 2a, the function *v*_*C*_(*t*) is plotted for a treatment plan consisting of one half hour of chemotherapy each day for 5 days followed by five days of rest, with this dosing cycle repeated 3 times. The time-course of the concentration of chemotherapy, *C*(*t*), resulting from this dose schedule is also plotted on the inset axis.

**Figure 2:**
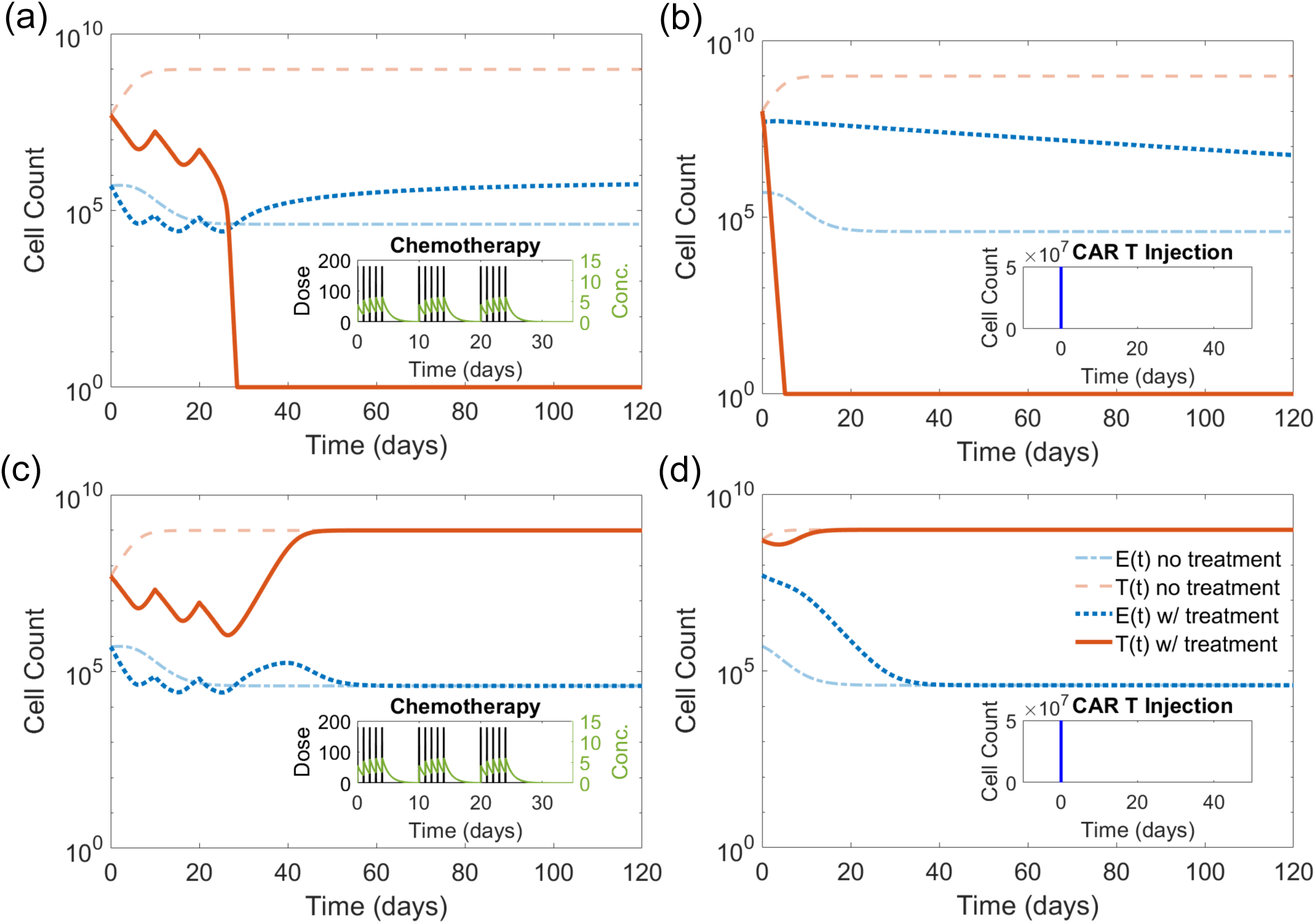
(a) A simulation demonstrating successful chemotherapy for an initial condition of (*T*_0_, *E*_0_) = (5 × 10^7^cells, 5 × 10^5^cells) with parameters from column 3 of Table 1. Without treatment, the patient trajectory would follow the dotted lines reaching the unhealthy outcome. However, intervening with 3 cycles of 5 days of chemotherapy at strength *S* = 175*µ*M/day alternating with 5 days rest (inset) results in tumor elimination and a healthy outcome. (b) A simulation demonstrating successful CAR T-cell therapy for an initial condition of (*T*_0_, *E*_0_) = (1 × 10^8^cells, 5 × 10^5^cells) using parameters from column 3 of Table 1. Without treatment, the patient trajectory would follow the dotted lines. However, an injection of 5 × 10^7^ CAR T-cells, plotted on the inset axis, results in tumor elimination and a healthy outcome. (c) The impact of the same chemotherapy plan on the same initial condition as Panel a, but instead using parameters from column 2 of Table 1. Again, without treatment, the patient trajectory reaches the unhealthy outcome. Under the treatment schedule depicted the, the tumor burden decreases initially but ultimately the trajectory still approaches the unhealthy patient outcome.(d) For the same patient parameters but an initial condition of (*T*_0_, *E*_0_) = (5 × 10^8^cells, 5 × 10^5^cells), the same injection of 5 × 10^7^ CAR T-cells now results in an unhealthy outcome

Following a method used to incorporate adoptive cellular therapy into previous models from the literature (de Pillis et al., 2006; Talkington et al., 2018), CAR T-cell injections of dose level *P* are modeled as a one-time increase in the number of effector cells, *v*_*E*_ (*t*), where

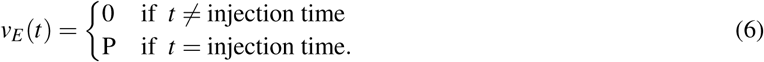

Altogether, the model system with treatment is

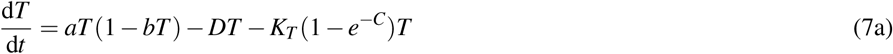

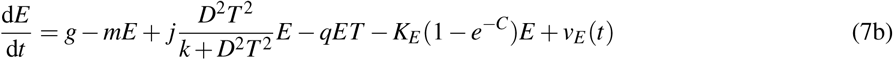

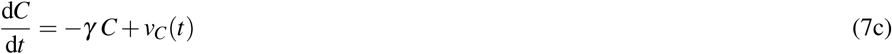

where again

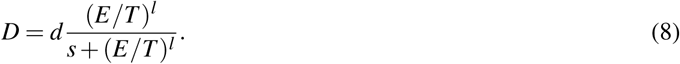

We model the impact of chemotherapy on the two cell populations by introducing saturating fractional cell-kill terms, *K*_*E*_ (1 − *e*^−*C*^)*E* and *K*_*T*_ (1 − *e*^−*C*^)*T*, to the equations for both tumor and effector cells in System (1a-b). These expressions saturate to the parameters *K*_*E*_ and *K*_*T*_ respectively as *C*(*t*) increases. Note that different types of chemotherapy can be considered by varying the associated parameters: *γ, K*_*T*_, and *K*_*E*_.

Model behaviors under individual chemotherapy and CAR T-cell therapy treatments are shown in Fig. 2. An acute dosage of chemotherapy is effective against some tumors (Fig. 2a), but milder dosages are not. Similarly, we verified that our model behaves as would be generally expected for CAR T-cell injections. In particular, we observed that perturbing the system with a large enough number of effector cells can shift the system so that it approaches the healthy equilibrium (Fig. 2b). We note the marked difference in the impact each type of therapy has on the effector cell population. Chemotherapy decreases the immune cell population, which then recovers over time. CAR T-cell injection boosts the initial effector cell population, which continues to increase for several days before slowly decreasing toward the equilibrium level.

Single-modality treatments, however, are often insufficient to achieve a desirable patient outcome, making combination treatments critical. A chemotherapy plan that is effective for one patient may still fail for other patients. For example, using the same chemotherapy plan that was successful in Fig. 2a against the same tumor burden, but with slightly different patient parameters results in an unhealthy patient outcome (Fig. 2c). Using a lower dosage of CAR T-cells may be necessary to reduce toxicity of treatment side effects to a tolerable level, but CAR T-cell injections alone can fail. If the same patient as Fig. 2b starts at a higher initial tumor burden of 5 × 10^8^ cells, the previously effective intervention initially slows tumor growth, but ultimately fails (Fig. 2d). The scenarios in Fig. 2c and 2d demonstrate cases where a single-treatment approach is ineffective. These limitations drive the need to combine treatment modalities in order to effectively combat cancer.

## 3 Evaluation of treatment plans

Standard treatment protocols incorporate a lymphodepleting preconditioning regimen which consists of a round of chemotherapy prior to CAR T-cell infusion. We tested realistic combinations of chemotherapy preconditioning and CAR T-cell doses by simulating treatment plans with parameter sets from several different cancer types and evaluating the outcomes. Here we discuss the scenarios considered and simulation results.

### 3.1 Treatment plans considered

We explored combinations of chemotherapy preconditioning and CAR T-cell doses from a range that encompasses the standard procedures for the two FDA approved CAR T-cell treatments, Yescarta (Kite, 2017) and Kymriah (Novartis, 2017). The guidelines for dosing CAR T-cells prescribe 0.2 − 5.0 × 10^6^ cells per kilogram of body weight up to a maximum of 2.5 × 10^8^ cells (Kite, 2017) or 6 × 10^8^ cells (Novartis, 2017). We consider injections of *P* = 1 × 10^7^ cells, *P* = 1 × 10^8^ cells, and *P* = 2.5 × 10^8^ cells referred to as dose level 1 (**DL1**), dose level 2 (**DL2**), and dose level 3 (**DL3**) respectively.

The standard chemotherapeutic preconditioning regimen for both drugs is a combination of fludarabine and cyclophosphamide administered over three to five days. We considered two chemotherapy regimens, either medium strength for one half hour each day for 5 days (**C5**), or high strength for one half hour each day for 3 days (**C3**). The strengths were chosen so that the peak concentration of chemotherapy during the five day regimen matched the value reported in a pharmacokinetic study of fludarabine (Ju et al., 2014), and the overall exposure in the two plans was equal (measured by the area under the chemotherapy concentration curve). Holding the administered dose (measured by the area under dosing curve, *ν*_*C*_(*t*)) constant between the two plans instead results in the same qualitative patient outcomes. In practice, lymphodepletion is followed by either a 3-day rest period for Yescarta (Kite, 2017) or 2-14 days rest for Kymriah (Novartis, 2017) before T-cell injection. For all combinations, we started by enforcing the minimum rest period of 2 days between the end of chemotherapy and the T-cell injection.

### 3.2 Patients

We considered four patient parameter sets in our evaluation. The first patient constitutes typical parameter values for diffuse large B-cell lymphoma (Rösch et al., 2016). Rösch et al. estimated a range of model parameters using data from randomized clinical trials in elderly patients. Our second patient parameter set is drawn from typical parameter values for B-cell chronic lymphocytic leukemia (Nanda et al., 2013). Nanda et al. analyzed existing data in the medical literature to determine ranges of values for model parameters and calibrated their model to clinical patient data. Currently CAR T-cell therapy has been approved for use with several blood cancers; however, numerous clinical trials are testing its potential for treatment of solid cancers as well, including melanoma (Brown and Mackall, 2019; Zhang et al., 2014). De Pillis et al. (2005) used data from two metastatic melanoma patients reported in a clinical trial to fit their model for mixed chemotherapy and immunotherapy treatment. We use their resulting parameter values for our last two patient parameter sets. The model assumptions are predicated on CAR T-cells effectively targeting tumor cells. Hence simulation results suggest possible outcomes once good target antigens have been identified for metastatic melanoma. The parameters we used for Patient 1-4 are listed in Table 1, and the details of parameter selection are discussed in Appendix B.

Partitioning the phase plane into the basins of attraction for the high-tumor and zero-tumor equilibria for each set of patient parameters provides intuition into how they will respond to treatment. The basins of attraction for Patient 1-4 are illustrated in Fig.3. If a trajectory is still in the unhealthy basin of attraction (shaded area) for a given patient after both chemotherapy and CAR T-cell injection, that treatment plan will not be effective. Because the “slope” of the boundary between the basins of attraction is less steep for Patients 3 and 4, CAR T-cell injection will be very effective regardless of preconditioning. On the other hand for Patient 2, the slope of the basin boundary is very steep so CAR T-cell therapy alone will not be effective. For Patient 1, we see that the basin boundary has an intermediate slope and so preconditioning with chemotherapy has the potential to make a big impact on the treatment outcome. The larger basin of attraction for the high-tumor equilibrium in Patients 1 and 2 results from differences in patient parameters. In particular, the carrying capacity for Patients 1 and 2 is on the order of 10^13^, whereas the carrying capacity for Patients 3 and 4 is on the order of 10^9^ because of the lower constraints on the number of tumor cells in hematological cancers compared to solid cancers.

**Figure 3:**
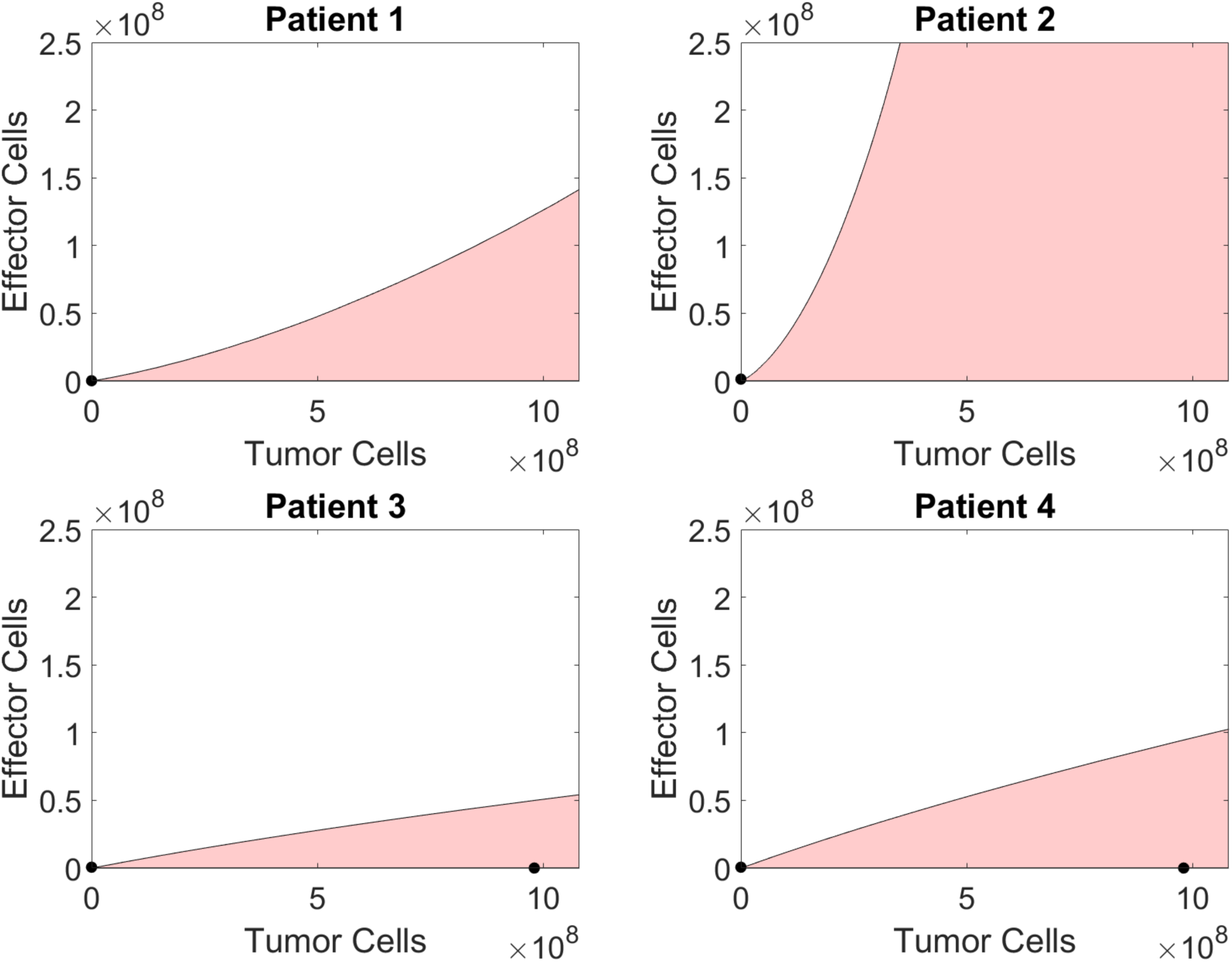
For Patients 1-4, the positive quadrant of the phase plane is partitioned into the basin of attraction for the zero-tumor equilibrium (white) and the high-tumor equilibrium (shaded). Stable equilibria are marked by circles. For Patients 1 and 2 the high-tumor equilibrium lies outside the figure

The initial conditions tested for each patient cover a range of tumor magnitudes. For Patient 3 and 4 the minimum value is close to the smallest detectable tumor size and the maximum is the carrying capacity for tumor cells in the absence of effector cells. The values considered were a low tumor burden (low TB) at 1 × 10^8^ tumor cells, medium tumor burden (medium TB) at 5 × 10^8^ tumor cells, and high tumor burden (high TB) at 9.8 × 10^8^ tumor cells (de Pillis et al., 2006). For Patient 1 and 2, we tested initial conditions an order of magnitude higher to reflect the higher tumor cell carrying capacity. The values considered for Patients 1-2 were a low tumor burden (low TB) at 1 × 10^9^ tumor cells, medium tumor burden (medium TB) at 5 × 10^9^ tumor cells, and high tumor burden (high TB) at 1 × 10^10^ tumor cells. The initial number of effector cells was set to 4 × 10^5^ cells, which is the average of the zero-tumor equilibrium values for all four patient parameter sets (Rösch et al., 2016). The estimated average number of T-cells activated against a given antigen is higher; however, cancer cells are notoriously hard for the immune system to recognize, so it is reasonable to consider a smaller initial immune response (Pandya et al., 2016).

### 3.3 Single treatment plan outcomes

First, we tested the CAR T-cell doses and lymphodepleting regimens one at a time to verify the impact on each tumor burden. In all scenarios, lymphodepleting chemotherapy alone resulted in tumor progression.

The results of CAR T-cell treatment without lymphodepletion are summarized in the first 3 columns of Fig. 4. For Patient 1 and 2, CAR T-cell therapy alone does not clear any of the initial tumor-burdens. For Patients 3 and 4, DL1 is insufficient, but DL 2 and above result in a healthy patient outcome in all scenarios. The results demonstrate that there are scenarios in which reasonable single-treatment plans were not sufficient to shift the patient initial condition into the healthy basin of attraction, highlighting the importance of combination treatment.

**Figure 4:**
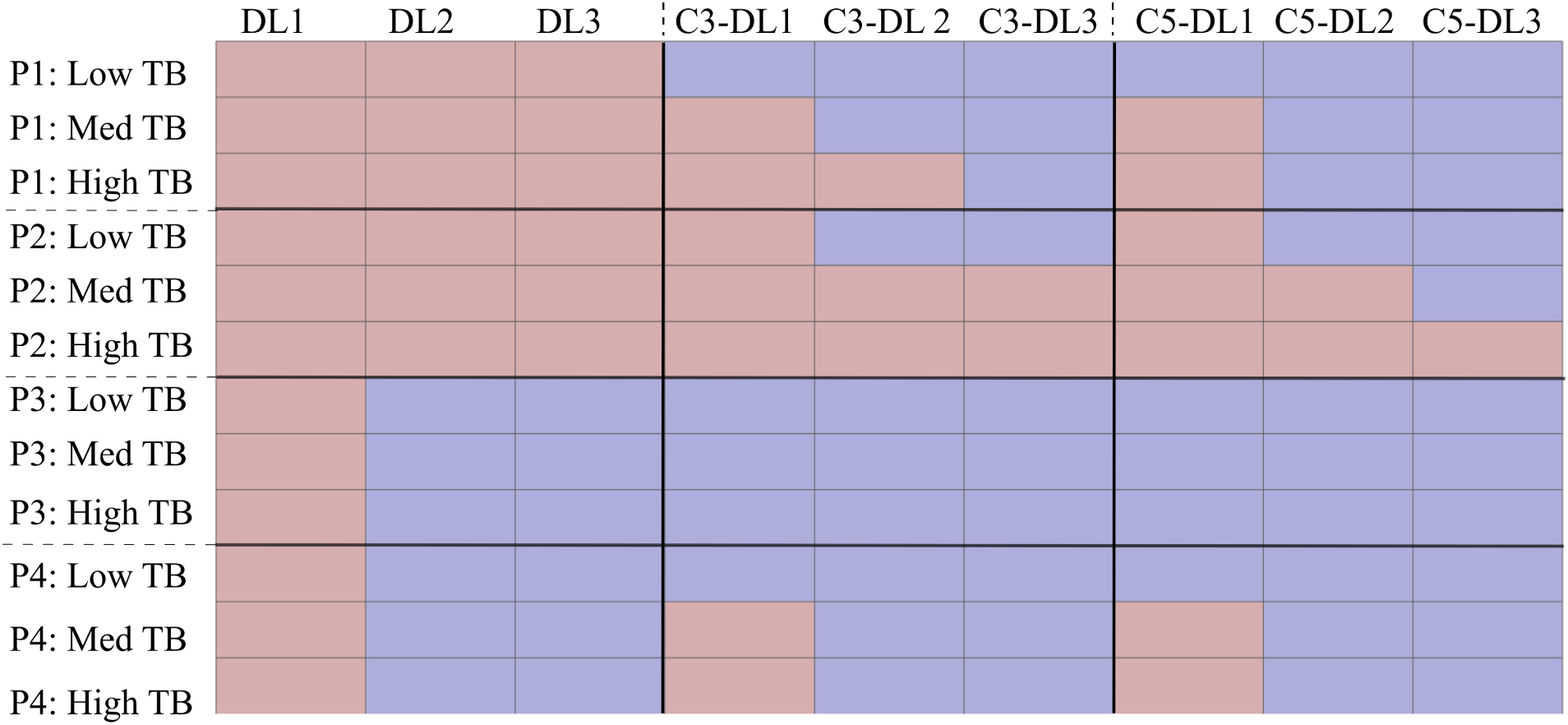
Treatment outcomes are recorded for Patients 1-4 according to the following key: DL1 = 1 × 10^7^ cells, DL2 = 1 × 10^8^ cells, DL3 = 2×.5 10^8^ cells, C3 is 3 days of chemotherapy at high strength, C5 is 5 days of chemotherapy at medium strength, a blue box indicates a healthy outcome, and a red box indicates an unhealthy outcome. For example, all plans expect DL1 alone were successful for Patient 3 with low TB. However, all plans without preconditioning failed for Patient 1

### 3.4 Combination treatment plan outcomes

After testing single-modality treatments, combination treatments were considered. The last 6 columns of Fig. 4 summarize the outcomes of combination treatments. Notably, for Patient 1 and 2, we observe conditions that cannot be treated by CAR T-cell injection alone, but are treatable with chemotherapy and CAR T-cell injection in combination. We also observe that preconditioning reduces the dosage of CAR T-cells necessary for effective treatment in many scenarios (Patient 1 all initial conditions, Patient 2 medium and low TB, and Patient 3 all conditions, and Patient 4 low TB). For the dose levels considered here, CAR T-cell injection and lymphodepleting chemotherapy were never effective for Patient 2 with high TB.

Between the fixed preconditioning regimens considered, the 3-day high-strength and the 5-day medium-strength chemotherapy dosing schedules often had similar outcomes. However, for Patient 1 high TB, combining 5-day chemotherapy with DL2 was effective in eliminating the tumor, but combining 3-day chemotherapy with the same dose of CAR T-cells was not. Similarly for Patient 2 medium TB combining 5-day chemotherapy with DL3 was effective in eliminating the tumor, but combining 3-day chemotherapy with the same doses of CAR T-cells was not. Plots of the time course of the trajectories under these two treatment plans for Patient 1 high TB are shown in Fig. 5. Without treatment the tumor-cell count climbs towards the carrying capacity and the effector-cell count drops, approaching the high-tumor equilibrium. Introducing 3 days of high strength chemotherapy reduces the tumor burden, but when chemotherapy ends and the CAR T-cell injection is administered on day 5, the tumor-cell count still climbs to carrying capacity (Fig. 5a). In contrast, 5 days of medium strength chemotherapy reduces the tumor burden enough so that after the CAR T-cell injection is administered on day 7 the tumor-cell count steadily declines to zero (Fig. 5b). Although the area under the chemotherapy concentration curve is the same in these two scenarios, spreading the concentration over 5 days instead of 3 dramatically changes the patient’s outcome.

**Figure 5:**
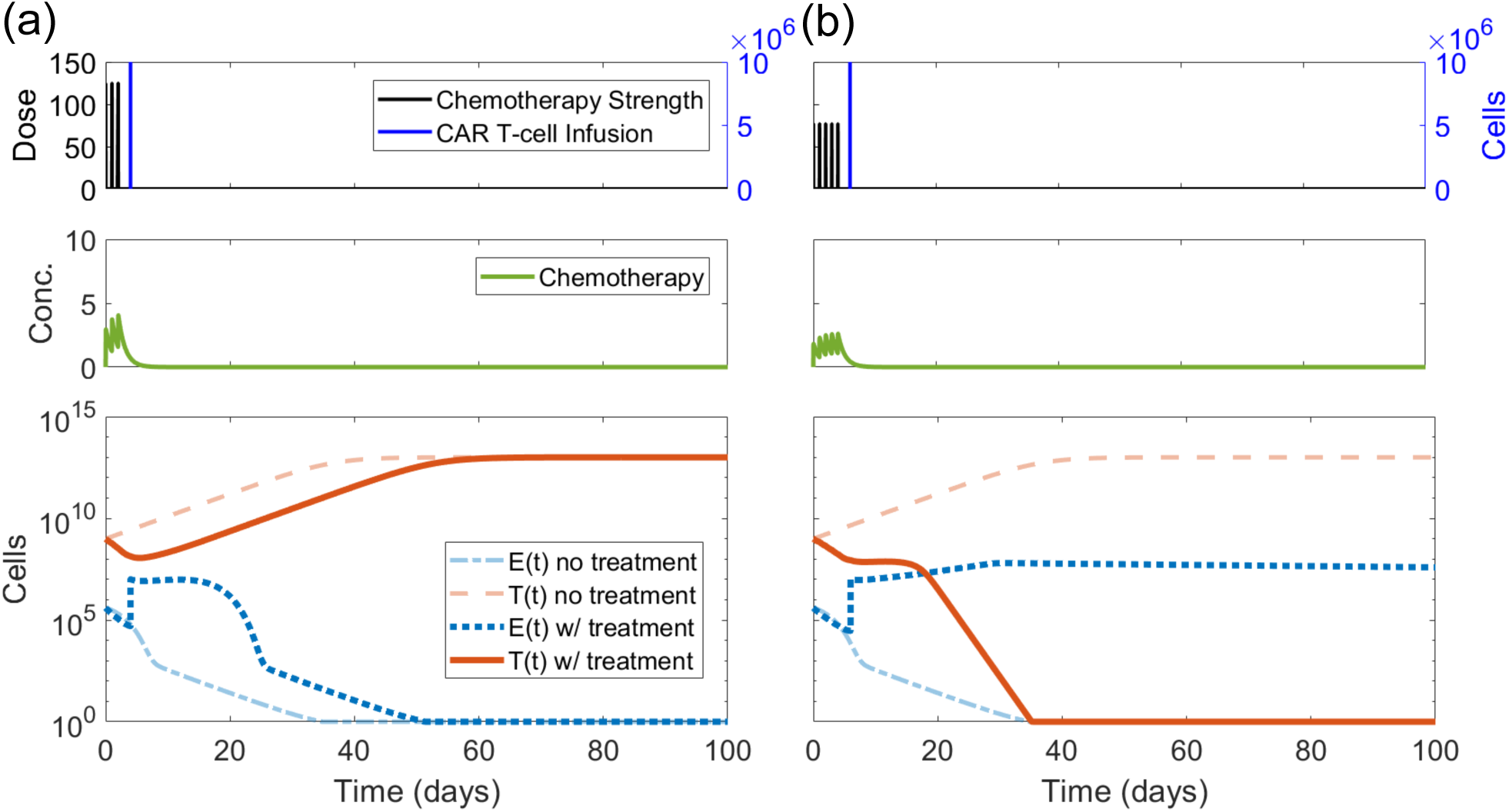
Numerical simulations of treatment of high TB for Patient 1. (a) Three days of high-strength chemotherapy are combined with a CAR T-cell injection of 1 × 10^8^ cells, producing an unhealthy outcome. (b) The 5-day medium strength chemotherapy plan is combined with the same CAR T- cell injection resulting in a successful patient outcome

Finally, our simulations showed that the rest period between the end of preconditioning and T-cell injection impacts patient outcome. For example, administering 3 days of high-strength chemotherapy, allowing two days of rest, and then administering an injection of CAR T-cells at DL1 is an effective intervention for Patient 3 high TB (Fig. 6a). However, if the rest period is extended to four days instead of two, the combination is no longer successful (Fig. 6b). During the extended rest period, the tumor-cell count climbs high enough that a CAR T-cell injection that was previously effective now fails. For each combination of chemotherapy and CAR T-cell therapy that succeeded with a 2-day rest period where the CAR T-cell dose alone failed, we determined the maximum possible rest period with which the combination was still effective. Over 50% of these scenarios failed if the rest period was 6 days or greater. Nearly 85% of the scenarios failed if the rest period was 12 days or longer. All of the scenarios failed if the rest period was longer than 16 days. This suggests that the length of the rest period between chemotherapeutic lymphodepletion and CAR T-cell injection can have important implications for patient outcome. As rest periods can range widely, from 2-14 days, under current guidelines this is an important aspect of treatment to investigate more thoroughly.

**Figure 6:**
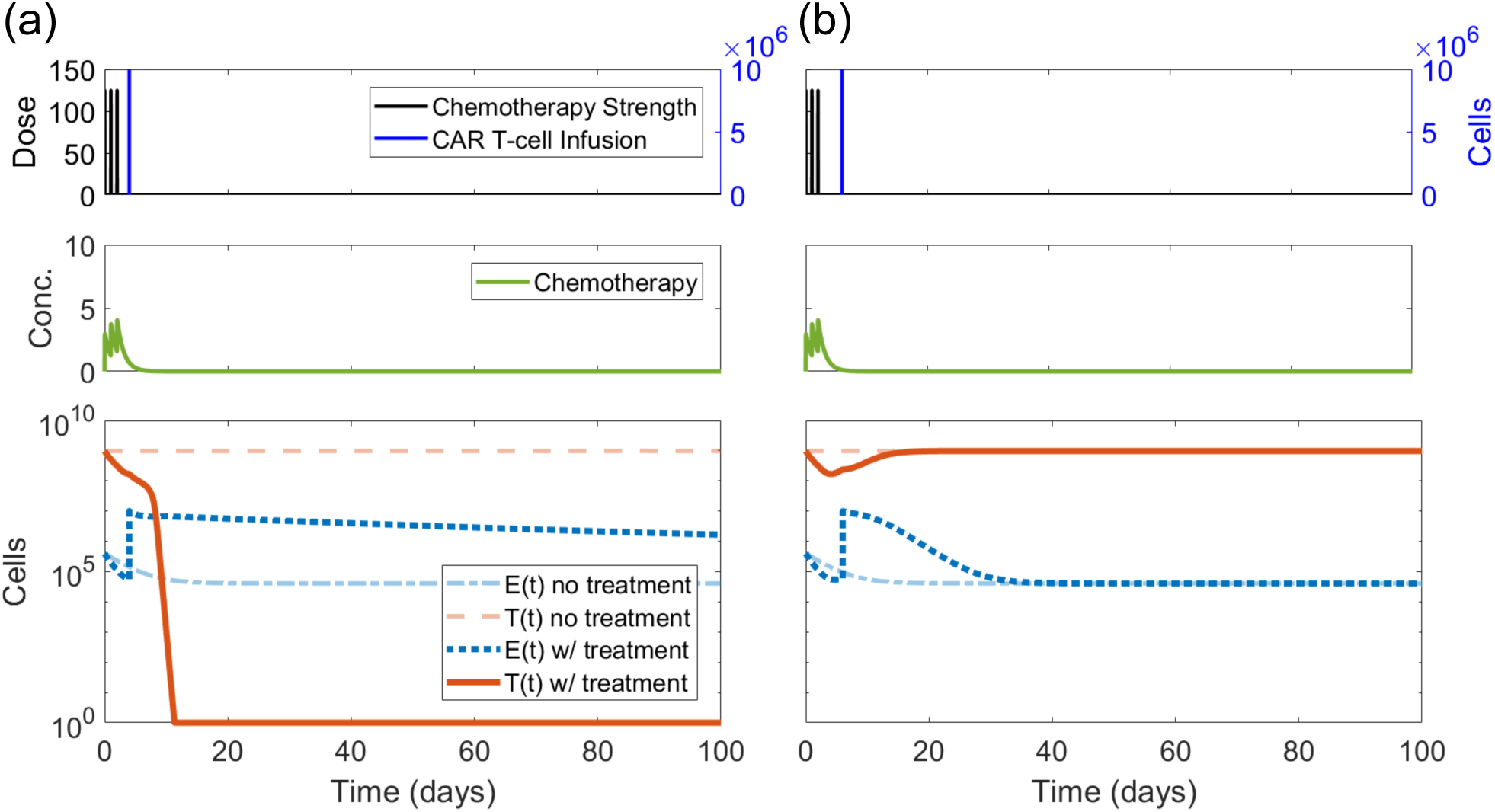
(a) Combining 3 days of high-strength chemotherapy and 1 × 10^7^ CAR T-cells sequentially with a 2-day rest period results in tumor elimination and a healthy outcome. (b) However combining the two treatments with a 4-day rest period in between allows the tumor to escape. Both simulations were run using parameters from Patient 3 and assuming high TB

### 3.5 Sensitivity Analysis

We performed a sensitivity analysis on the model to identify which parameters have the largest impact on the effectiveness of CAR T-cell injections. For each patient, we calculated the precise threshold of effector cells above which a patient initial condition will move towards the healthy, tumor-free equilibrium when the number of tumor cells is at the smaller of the two high TB, 9.8 × 10^8^ cells. We call this number of effector cells the high-tumor threshold, because any treatment plan which injects a dose of CAR T-cells exceeding this quantity will be successful (in theory) against tumor burdens up to and including the high TB (Fig. 7).

**Figure 7:**
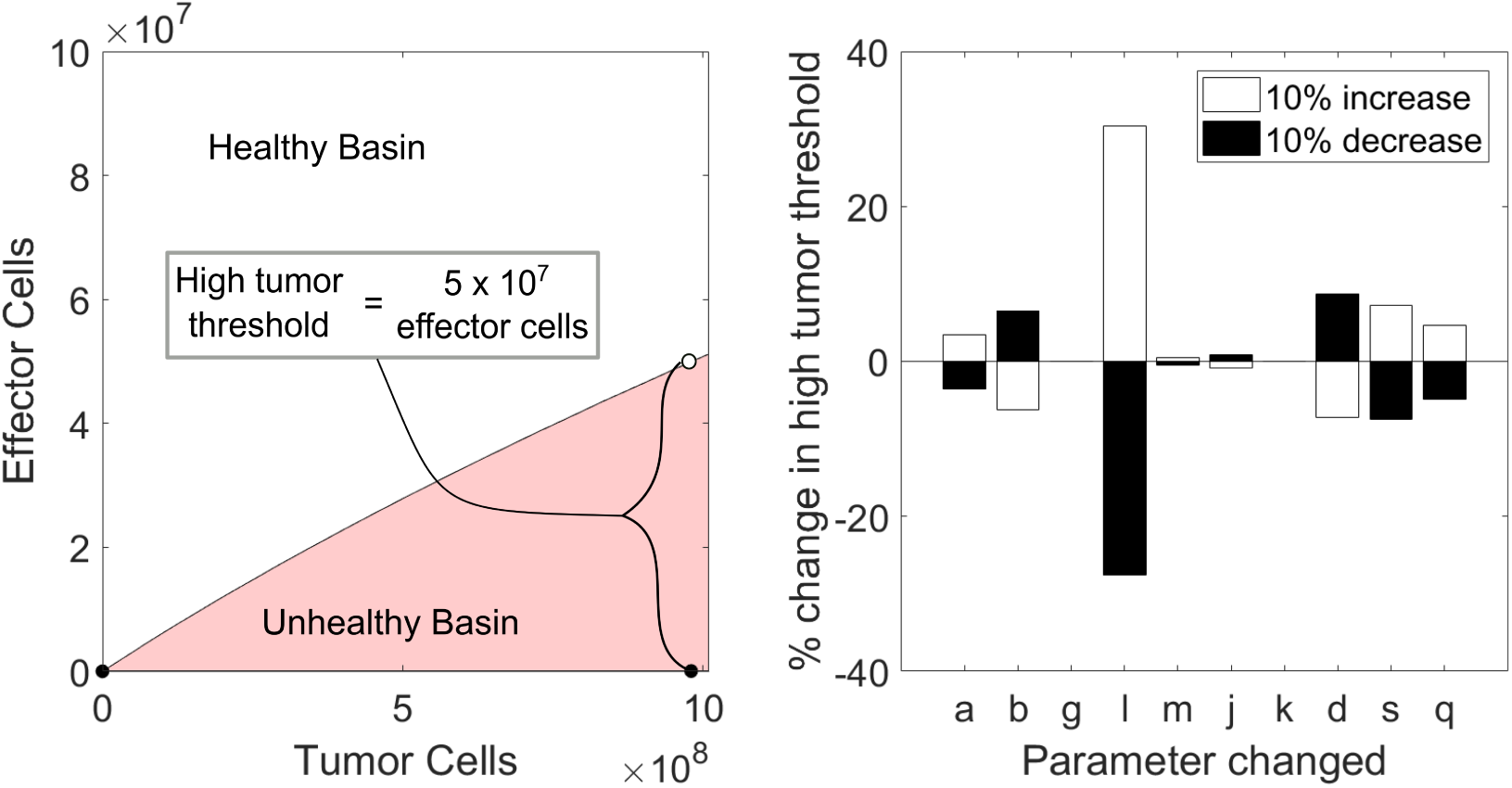
We define the high tumor threshold as the height of the boundary between the healthy basin and the unhealthy basin at high tumor burden. That is, the minimum number of effector cells needed to contain a disease burden of 9.8 × 10^8^ tumor cells. Sensitivity analysis shows that the high tumor threshold is most sensitive to the tumor-cell lysis rate exponent, *l*. These values were computed using the Patient 3 parameter set.

For the sensitivity analysis we varied each parameter up and down by 10% while holding the other parameters constant and calculated the resulting change in the high-tumor threshold. (See Appendix C for details of sensitivity analysis.) The exponent in the tumor-cell lysis rate, *l* ∈ [1, 2], has the largest impact on the high-tumor threshold (Fig. 7). The fact that a 10% change in *l* can result in a 30% shift in the threshold indicates that small changes in CAR T-cell effectiveness against tumor cells will affect clinical outcomes. The next most influential parameters are *d* and *s*, which also define the shape of the ratio-dependent cell-lysis term, with *d* setting the maximum lysis rate and *s* affecting how steeply the fractional term saturates to *d*. Again, these parameters relate to the effectiveness of CAR T-cell anti-tumor activity.

In contrast, changes in the parameters defining the base recruitment rate of effectors cells, *g*, the maximum recruitment rate of effector cells by tumor-cell lysis, *j*, the steepness of the effector cell recruitment curve, *k*, and the death rate of effector cells, *m*, had little effect on the high-tumor threshold. This suggests that incremental changes in the effectiveness of CAR T-cells may reduce the dosage required for successful therapy more than small changes in the recruitment and proliferation of CAR T-cells.

## 4 Discussion and conclusion

We have developed and analyzed a mathematical model consisting of a system of ordinary differential equations that can be used to test combinations of preconditioning chemotherapy regimens and CAR T-cell doses. We also found that under biologically relevant parameter values the system has two stable equilibrium points, one “healthy,” tumor-free equilibrium and one “unhealthy,” high-tumor equilibrium. The basins of attraction for these two equilibria were determined numerically. The variation between the shapes of the basins of attraction for different parameter sets reflect the fact that responses to treatment will not be uniform across patients.

We tested treatment plans that adhere to standard CAR T-cell therapy protocols on patient parameter sets from several cancer types. Our goal was not to propose any novel therapy plans or unrealistic dosing schedules, but rather to uncover refinements that are attainable within current practice and that merit additional attention. We observed a variety of outcomes supporting three main conclusions:

First, CAR T-cell injections will not be effective if a patient’s tumor burden is too high. Exactly what is meant by “too high” is patient specific, but can be quantified by characterizing the boundary between the basin of attraction for the healthy versus unhealthy equilibria. With this in mind, for many patients effective pre-conditioning is crucial to reduce tumor burden below the threshold before injecting CAR T-cells. One model parameter closely associated with the slope of the boundary is the exponent of the tumor-cell lysis rate, *l*, which indicates how efficient effector cells are at killing tumor cells. This observation suggests that small changes in the effectiveness of CAR T-cells can have a large impact on the dosage required for successful adoptive cellular therapy.

Second, appropriate chemotherapeutic lymphodepletion can reduce the CAR T-cell dosage necessary for successful treatment. This was observed in treatment scenarios for all four patients considered. Lower tumor burden at the time of injection and lower CAR T-cell doses are associated with milder side effects, which implies that selecting the appropriate lymphodepletion plan can make CAR T-cell therapy safer.

Finally, the recovery period between when preconditioning chemotherapy ends and a CAR T-cell injection is administered matters. If the recovery period is too long, the benefits of lymphodepletion may be lost. This finding in particular warrants further investigation as under current practice rest period can vary from as low as 2 up to as high as 14 days. Ideally, model parameters should be tuned to match a patient’s tumor growth rate in order to suggest how strictly the rest period should be limited.

CAR T-cells have transformed treatment of hematological cancers and show enormous potential for further innovation and application. One challenge facing this technology is the ubiquity of severe side-effects. Standard treatment protocol includes a chemotherapeutic preconditioning regimen, yet the optimal combination of chemotherapy and CAR T-cells to minimize side-effects while maintaining efficacy has not been determined. Addressing the question of how these two forms of treatment interact for any given patient with a simple mathematical model takes an inexpensive first step towards informing further investigations.

## Appendix A: Model Analysis

To begin our analysis, we first non-dimensionalize Sys. (2) as follows. Let *x* = *bT, y* = *aE/g, D*^*^ = *D/a*, and *t*^*^ = *at*. This results in seven non-dimensional parameters,

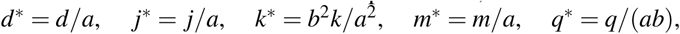

and *s*^*^ = *s*(*a/*(*gb*))^*l*^. The parameter *l* is nondimensional in the original system, and so remains unchanged. Dropping stars for notational simplicity, the non-dimensionalized system without treatment is given by

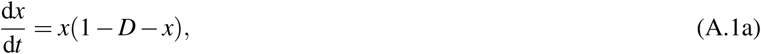

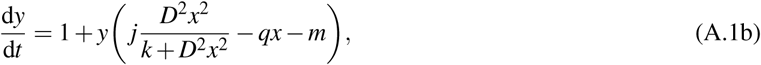

where *D* is a new dimensionless ratio term

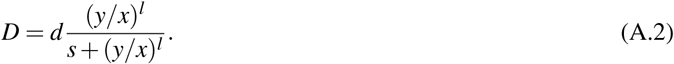

Next, we find the equilibria of the system, which occur at the intersections of the tumor and effector-cell nullclines, where 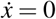 and 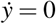, respectively. From Eq. (A.1a) we see that the tumor cell nullclines are *x* = 0 and *x* = 1 − *D*. For the effector cell nullcline, we set the right hand side of Eq. (A.1b) equal to zero and find that 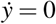 when

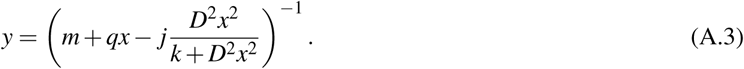

We will refer to the intersection between the tumor nullcline *x* = 0 and the effector cell nullcline, Eq. (A.3), as the zero-tumor or tumor-free equilibrium. Equilibria that occur at the intersections between the tumor nullcline *x* = 1 − *D* and the effector cell nullcline, Eq. (A.3), are referred to as nonzero-tumor equilibria or interior equilibria.

In order to determine the stability of the steady states, we apply linear stability analysis. The linearized system at point (*x, y*) is summarized by the Jacobian, *H*, of system (A.1) evaluated at (*x, y*). Let

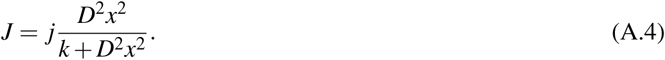

Then the Jacobian of Eq. (A.1) evaluated at that point (*x, y*) is,

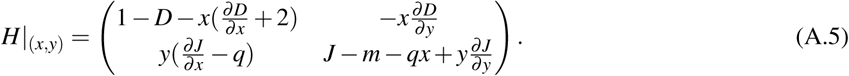

Let

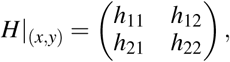

so that substituting the partial derivatives into Eq. (A.5) gives

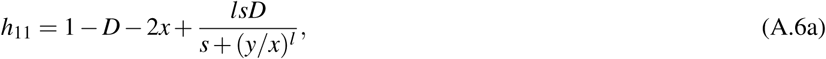

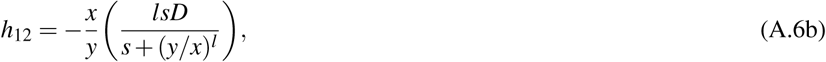

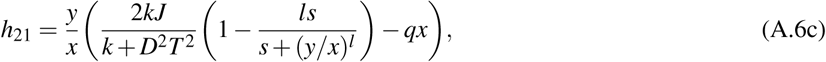

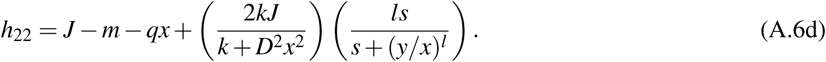

The simplest steady-state to characterize is the tumor-free equilibrium. Evaluating the effector cell nullcline at *x* = 0 yields *y* = 1*/m*. Hence, the tumor-free equilibrium occurs at (*x*_0_, *y*_0_) = (0, 1*/m*). Linearizing about this point, the Jacobian is

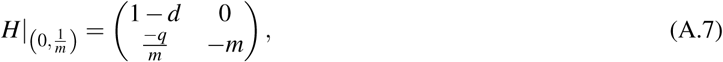

which has eigenvalues *λ*_1_ = 1 − *d* and *λ*_2_ = − *m*. The parameter values *d* and *m* will always be positive real numbers. It follows that *λ*_1_ and *λ*_2_ are always real and positive when *d* > 1. For the biologically relevant cases we considered, the parameter *d* > 1, so this condition is satisifed. Hence the equilibrium point will be a stable node, which aligns with the results reported by de Pillis et al. for their 2006 model. The stability of the tumor-free equilibrium reflects the idea that, once activated, the immune system can contain a tumor if it is small enough.

Now we turn our attention to the interior equilibria. For these equilibria

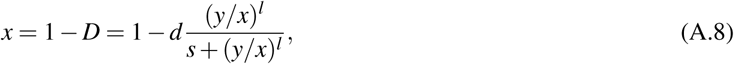

so we can solve for *y* as a function of *x* along the tumor nullcline,

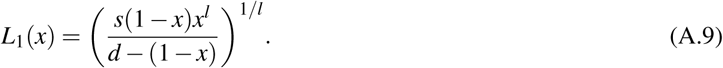

Substituting *x* = 1 − *D* into the effector cell nullcline and considering *y* a function of *x* yields

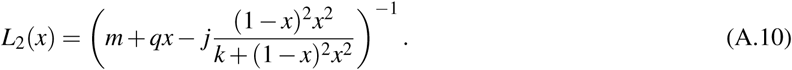

The nonzero-tumor equilibrium points are the intersections of Eq. (A.9) and Eq. (A.10) for each set of parameters. See Fig. 1a for a plot of *L*_1_(*x*) and *L*_2_(*x*) for the Patient 3 parameters with equilibria marked. For reasonable parameter ranges, reported in Table 1, we observed that the model has two positive nonzero-tumor equilibrium points because the parameter *s* is large enough that *L*_1_(*x*) exceeds *L*_2_(*x*) at some point on the interval *x* ∈ (0, 1), but *L*_1_(0) = 0 < *L*_2_(0) and *L*_1_(1) = 0 < *L*_2_(1).

The first interior equilibrium occurs at a large number of tumor cells, 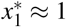, and a small number of effector cells, 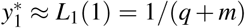. The second interior equilibrium occurs at a small number of tumor cells and a moderate number of effector cells. Although it is not possible to find exact, closed-form expressions for the interior equilibria because it involves finding the roots of a quintic polynomial, we can use the Routh-Hurwitz criterion to find conditions under which the high-tumor equilibrium is stable (Wiggins, 2003, p. 13). First, we use the nullclines to simplify the Jacobian. Because we are at the intersection of *L*_1_ and *L*_2_, it must hold that *D* = 1 − *x* and *J* − *m*− *qx* = − 1*/y* so the Jacobian has entries

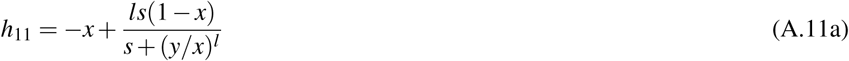

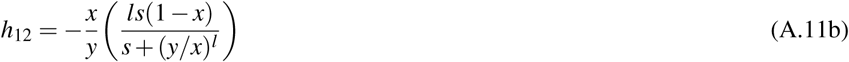

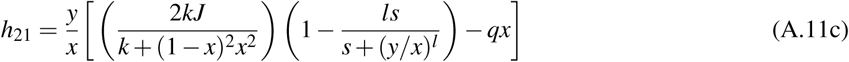

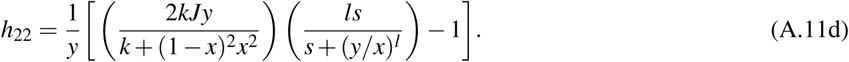

Now consider the trace of the Jacobian,

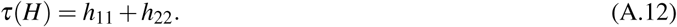

From Eq. (A.11a), we can see that when *x* ≈ 1 it follows that that *h*_11_ ≈ −1. Observe that because *J* ≤ *j* and both *k/*(*k* + (1 −*x*)^2^*x*^2^) and *s/*(*s* + (*y/x*)^*l*^) are less than 1, we also have an upper bound on *h*_22_. It must be that

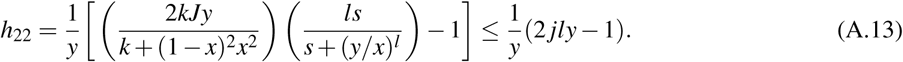

Hence for *y* < 1*/*(2 *jl*), entry *h*_22_ is negative. Thus when *x* ≈ 1 and *y* < 1*/*(2 *jl*), both *h*_11_ and *h*_22_ will be negative, so the trace must be negative.

Next consider the determinant of the Jacobian

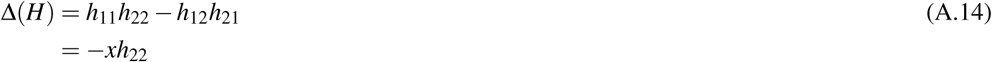

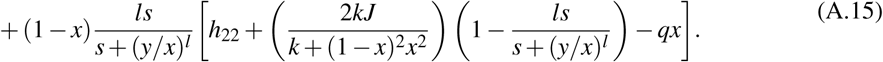

When *x* ≈ 1, then (1 −*x*) is approximately zero which leaves

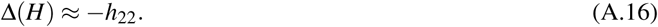

As established in Eq. (A.13), when *y* < 1*/*(2 *jl*) it follows that *h*_22_ < 0, so Eq. (A.16) indicates that under this same condition Δ(*H*) > 0. Thus the trace, *τ*(*H*), is negative and the determinant, Δ(*H*), is positive when *x* ≈ 1 and *y* < 1*/*(2 *jl*). These conditions are satisfied by the high-tumor equilibrium, so it follows by the Routh-Hurwitz criterion that the high-tumor steady state is stable.

In our numerical simulations, we observed that the low-tumor steady state occurs for a moderate number of effector cells and a small number of tumor cells resulting in a Jacobian with the signs of each entry as follows

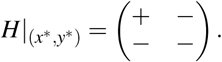

The first entry, *h*_11_, has the opposite sign from the high-tumor equilibrium because now 1 − *x*^*^ > *x*^*^. In this case, the determinant is negative, so the equilibrium is a saddle point, hence unstable. The tumor burden for this low-tumor equilibrium was on the order of 10^7^ cells or fewer for the range of parameters considered. For context, a 1 cm^3^ tumor is estimated to contain 10^8^ − 10^9^ (Del Monte 2009). Because of its relatively small size if the low-tumor equilibrium were stable, it could be considered an alternate healthy outcome in which the disease is contained but not eliminated.

The first quadrant is an invariant set of system A.1. In order for a trajectory to leave the first quadrant, it would have to cross an axis, which requires 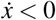 somewhere along the *x* = 0 axis or 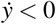 somewhere along the *y* = 0 axis. However, when *x* = 0 it is clear from Eq. (A.1a) that 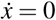. From Eq. (A.1b), we can also see that when *y* = 0, it follows that 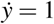, which is positive.

## Appendix B: Parameter Selection

We selected parameters based on previous models in the literature. The metastatic melanoma parameter sets came directly from the parameter values for Patients 9 and 10 in the governing equations for tumor cells and CD8^+^ T-cells reported by de Pillis et al. (2005). However to counteract the effect of eliminating the other immune cell types (NK cells and circulating lymphocytes) that stimulate CD8^+^T cells, the base recruitment rate of effector cells from Kuznetsov et al. (1994) was taken to be the base recruitment rate of effector cells for Patients 3 and 4.

We set the parameter values for our diffuse large B-cell lymphoma patient to the midpoint of the ranges reported by Rösch et al. (2016) when there was an analogous term. We could use most parameters directly from Rösch et al. (2016). However, the parameters involved in the novel tumor-cell lysis term introduced by de Pillis et al. did not correspond directly to parameters in Rösch’s model. In formulating their model, Rösch et al. mention that effector cells will be kill tumor cells at a lower rate in lymphoma than in leukemia, so we chose to take *d* to be 3*/*4 of the *d* value for our leukemia patient. We set the remaining two parameters, *s* and *l*, to the average of the two patients considered in the model from de Pillis et al. (2005).

The chronic lymphocytic leukemia patient parameter values were set to the midpoint of the ranges reported by Nanda et al. (2013) when there was an analogous term, adjusting for differences in units. The Nanda et al. model assumed exponential tumor growth, rather than logistic so we set *b* = 10^−13^ because this choice allows tumor cells to exhibit essentially exponential growth in the absence of an immune response. Nanda et al. used a mass-action term to model the decrease in tumor cells due to contact with effector cells, so we set the saturating value of the tumor-cell lysis rate, *d*, to be the product of the mass action kill rate from Nanda et al. (2013) and the maximum dose of CAR T-cells considered. We set the final parameters involved in the novel interaction term (*l* and *s*) to the average of the two patients considered in the model from de Pillis et al. (2005).

The kill parameters relating to chemotherapy were taken from de Pillis et al. (2006), since the patients in the metastatic melanoma study were also treated with fludarabine. Accordingly, the fractional cell kills were set to *K*_*T*_ = 0.9 and *K*_*E*_ = 0.6 which assumes that chemotherapy is one log-kill and that it kills a larger fraction of tumor cells than host cells. The decay rate *γ* can be calculated from the half-life of a substance, *τ*, by *γ* = ln(2)*/τ*. The choice of strength for chemotherapy regimens was chosen by working backwards to see what values resulted in reasonable concentrations of fludarabine building up in the system, where “reasonable” concentrations matched those reported by Ju et al. in a pharmacokinetic study of the drug (2014). For 25 mg of fludarabine administered over one half hour each day for 5 days, Ju et al. reported a peak concentration of *C*_*max*_ = 1, 222(668 − 1, 732)ng/mL = 3.34 *µ*M. We found that a dose strength of *S* = 77 *µ*Mday^−1^ achieved this *C*_*max*_ in our simulations, so this was used as medium strength. The high strength was set to be *S* = 125 *µ*Mday^−1^ in order to achieve the same area under the concentration curve with only 3 days of injections. If we chose to instead match the dose administered (area under *ν*_*C*_(*t*)) between the two plans, the patient outcomes were qualitatively similar.

Note that while the treatment parameters are well-founded in pharmacokinetic data, the patient parameters have been drawn from previous theoretical studies for different cancer types. Some have been fit to patient data, but not all. In order to strengthen the impact of any suggestions made by this model it remains to calibrate parameters model to data from previous CAR T cell therapy patients and validate the model by evaluating its predictive performance on untrained data.

## Appendix C: Numerical Simulations and Sensitivity Analysis

We implemented numerical simulations with parameters for Patient 1-4 listed in Table 1 using a combination of 4th order Runge-Kutta integration schemes in MATLAB. During chemotherapy, we enforced a fixed step size of d*t* = 0.0208 corresponding to approximately one half hour or 1*/*48 of a day using ode4(). The remainder of each simulation was completed with ode45(), which dynamically chooses the time-step.

To discover which components of the model contribute most significantly to the effectiveness of CAR T-cell injections, we performed a numerical sensitivity analysis assessing how the model parameters impact the shape of the separatrix between the basin of attraction for the high-tumor equilibrium and the basin of attraction for the tumor-free equilibrium.

As a baseline for Patients 1,3 and 4, we calculated the precise threshold of effector cells above which a patient initial condition will move towards the healthy, tumor-free equilibrium when the number of tumor cells is at high TB, or 9.8 × 10^8^ cells. We call this number of effector cells the high-tumor threshold, because any treatment plan which injects a dose of CAR T-cells exceeding this quantity will be successful (in theory) against tumor burdens up to and including the high TB (Fig. 7). Then each parameter was increased by 10% while holding all other parameter values constant. The relative change in the high-tumor threshold was computed by subtracting the unperturbed high-tumor threshold from the perturbed high-tumor threshold and dividing the the difference by the unperturbed threshold. The same procedure was followed for a 10% decrease in each parameter. The high-tumor threshold with unperturbed parameters is essentially infinite for Patient 2, as can be inferred from Fig. 3, so this analysis could not be carried out.

The parameter which has the largest impact on the high-tumor threshold is the exponent in the tumor-cell lysis rate, *l* (Fig. 7). The parameter *l* is between 1 and 2 for all patient parameters considered and it encodes how the lysis rate depends on the ratio of effector/tumor cells. The fact that a 10% change in *l* can result in a 30% shift in the threshold indicates that small changes in CAR T-cell effectiveness will affect clinical outcomes. The next most influential parameters are *d* and *s*, which also define the shape of the ratio-dependent cell-lysis term, with *d* setting the maximum lysis rate and *s* affecting how steeply the fractional term saturates to *d*. Again, these parameters relate to the effectiveness of CAR T-cell activity.

## Acknowledgements

The authors would like to thank Kelsey Marcinko and Benjamin Liu for their helpful feedback during the writing process.

